# SARS-CoV2 quantification using RT-dPCR: a faster and safer alternative to assist viral genomic copies assessment using RT-qPCR

**DOI:** 10.1101/2020.05.01.072728

**Authors:** Paula Rodrigues de Almeida, Meriane Demoliner, Ana Karolina Antunes Eisen, Fágner Henrique Heldt, Alana Witt Hansen, Karoline Schallenberger, Juliane Deise Fleck, Fernando Rosado Spilki

**Affiliations:** Molecular Microbiology Laboratory, Feevale University

## Abstract

In this study, serial dilutions of SARS-CoV 2 RNA extract were tested using RT-dPCR using three different primer-probe assays aiming SARS-CoV 2 nucleocapsid coding region. Narrower confidence intervals, indicating high quantification precision were obtained in 100 and 1000-fold serial dilution and RT-dPCR results were equivalent between different assays in the same dilution. High accuracy of this test allowed conclusions regarding the ability of this technique to evaluate precisely the amount of genomic copies present in a sample. We believe that this fast and safe method can assist other researchers in titration of SARS-CoV2 controls used in RT-qPCR without the need of virus isolation.

## Introduction

Human coronaviruses (CoV) were often associated to mild respiratory diseases such as winter colds. In 2003, an outbreak of a severe acute respiratory disease (SARS) caused by CoV 1 changed the point of view of mankind about this virus (Holmes, 2003). In January of 2020, a new CoV (SARS-CoV2) emerges in Hubei province, China, causing severe respiratory disease, affecting over 80 thousand people in that region. Currently, the disease is known as COVID-19 (CoV disease 2019), the infection causes variable degree of severity, usually severe, seldom fatal in individuals with other diseases and the elderly, with case-fatality rate between 8% and 15% in this age in China. High transmissibility is the main difference between SARS-CoV2 and other CoV genotypes affecting humans (Wu & McGoogan, 2020, Velavan & Meyer, 2020). The high rate of transmissibility has impaired the local control of SARS-CoV2 in China, allowing it to spread to Europe and Americas. In Italy and Spain, it has been especially severe, reaching higher case-fatality and causing a collapse of official health services (Lazzerini & Putoto, 2020). SARS-CoV2 infections are now in acceleration phase in the Americas, with over 200.000 cases in the US, and over 1 million cases around the world.

Real-time information is among the most important factors helping to control local spread of COVID-19. Additional information about viral titration methods in different samples are necessary to estimate viral pathogenicity, to assess treatment success, estimate viral degree of transmissibility and viral survival in different surfaces and situations.

In order to titrate Sars-CoV2 via classical viral culture methods, a Biosecurity Level 3 (BSL3) facility is required, moreover, these methods are time consuming, and impose a higher risk of infection for people involved. Here, we present a fast, accurate and simple method of viral titration using QuantStudio 3D® microchip based RT-dPCR to titrate SARS-CoV2 genomic copies from controls to be used in RT-qPCR assays for diagnosis and research purposes.

## Material and Methods

RNA was extracted with MagMax™ RNA isolation kit and cDNA was synthesized with Promega GoScriptTM according to manufacturer’s instructions. Next, cDNA was diluted in a 10-fold based series from 10^−6^ to 10^−1^ to attempt ideal target copy numbers in the final reaction.

Optimal target concentration is essential for an accurate quantification through RT-dPCR (Majumdar et al., 2015). A dilution that results in approximately 200 to 2000 target copies in the final reaction usually presents better precision values. To achieve this titration of target copies, a series of RT-dPCR were attempted in six serial 10-fold dilutions.

Digital RT-dPCR was performed in a QuantStudio 3D™ System (Applied Biosystems™), using QuantStudio™ 3D Digital PCR Master Mix v2 dPCR mastermix and QuantStudio™ 3D 20K v2 chips. Primers and probe targeting SARS-CoV2 nucleocapsid 1, 2 and 3, described by (FDA) were used in RT-dPCR reactions. Cycling conditions were as follows: in 10 min at 96°C, followed by 39 cycles of 30 sec at 60°C and 2 min at 98°C, and a final step of 60°C per 2 min followed by maintenance at 4°C. V2 chips were read in the QuantStudio 3D instrument and the results were interpreted in the dPCR AnalysisSuite™ app in the Thermo Fisher Connect ™ Dashboard. Results with precision values below 5% were selected to estimate quantity of SARS-CoV2 genomic copies based on RT-dPCR.

## Results and Discussion

Ideal precision values were achieved in all three assays in dilutions of 1:10^−2^ and 1:10^−3^ahighlighted in Table 1.

**Table 1.** RT-dPCR results of 10-fold serial dilution of SARS-CoV2 control using assays for three targets in the Nucleocapsid gene. Selected samples are highlighted in bold.

| Dilution | Target | Average (copies/μL) | Interval (copies/μL) | Precision |
| --- | --- | --- | --- | --- |
| SARS-CoV$\cdot 10^{-6}$ | N1 | NA | NA | NA |
| SARS-CoV$\cdot 10^{-5}$ | N1 | 3,175 | 2,319 -- 4,345 | 36.88% |
| SARS-CoV$\cdot 10^{-4}$ | N1 | 31,655 | 28,506 -- 35,151 | 11.05% |
| **SARS-CoV$\cdot 10^{-3}$** | **N1** | **320,23** | **309.81 -- 331.01** | **3.37%** |
| **SARS-CoV$\cdot 10^{-2}$** | **N1** | **2511,8** | **2465.4 -- 2559.1** | **1.88%** |
| SARS-CoV$\cdot 10^{-1}$ | N1 | 9883,4 | 9103.7 -- 10730 | 8.56% |
| SARS-CoV$\cdot 10^{-6}$ | N2 | 1087 | 0.602 -- 1.963 | 80.57% |
| SARS-CoV$\cdot 10^{-5}$ | N2 | 3,69 | 2.685 -- 5.071 | 37.44% |
| SARS-CoV$\cdot 10^{-4}$ | N2 | 35,51 | 32.129 -- 39.247 | 10.52% |
| **SARS-CoV$\cdot 10^{-3}$** | **N2** | **353,51** | **342.13 -- 365.28** | **3.33%** |
| **SARS-CoV$\cdot 10^{-2}$** | **N2** | **2466,6** | **2417.7 -- 2516.3** | **2.02%** |
| SARS-CoV$\cdot 10^{-1}$ | N2 | NA | NA | NA |
| SARS-CoV$\cdot 10^{-6}$ | N3 | 0,832 | 0.433 -- 1.599 | 92.19% |
| SARS-CoV$\cdot 10^{-5}$ | N3 | 3,344 | 2.401 -- 4.658 | 39.28% |
| SARS-CoV$\cdot 10^{-4}$ | N3 | 30,958 | 27.899 -- 34.353 | 10.96% |
| **SARS-CoV$\cdot 10^{-3}$** | **N3** | **344,74** | **333.47 -- 356.39** | **3.38%** |
| **SARS-CoV$\cdot 10^{-2}$** | **N3** | **2531,9** | **2480.9 -- 2584** | **2.06%** |
| SARS-CoV$\cdot 10^{-1}$ | N3 | NA | NA | NA |

Samples highlighted in Table 1 were selected to estimate genomic copies of SARS-CoV2 in the positive control stock to be used in future RT-qPCR assays. These samples presented a low precision value. Precision value is calculated based on the confidence interval estimated for the sample. Target titration through digital PCR relies on Poison distribution using chips with microscopic partitions where individual PCR reaction occurs with a probe based assay, allowing a quantification based on negative to positive rates among these partitions. Chips used in this experiment have 20.000 partitions, in general, 20% of negative partitions results in good precision values. Precision of RT-dPCR quantification indicates how reliable is the result achieved in a reaction, it is directly related to confidence interval, with a narrower confidence interval resulting in desirable precision values and consistent target quantification results. Precision is dependent on target concentration, and ideal precision can be achieved by serial dilutions. Ideal concentrations are those with lower precision values and narrower confidence intervals (Majumdar, et al., 2015).

In this experiment three primer-probe assays were applied to titrate viral genomic copies per microliter. It is possible to observe very close quantification and precision results for the three probes at the same concentration. Distribution of reactions in the chips were also very similar, as shown in Figure 1. These results indicate that these primer-probe assays are suitable for SARS-CoV2 quantification through RT-dPCR. All three assays target the same gene, but in different locations and the similarity of the results in two different concentrations and 3 different probes reinforce the reliability of this technique in titration of genomic copies without need of excessive direct viral manipulation and time consuming techniques. The limit of detection reported for these assays was of 6,25 copies/µL (FDA). Control samples in diluted in 1:10−5 presented an estimated value of approximately 3 copies per µL, however here cDNA was diluted in water, moreover, these samples were above the cutoff value established for interpretation of reliable quantification assays.

**Figure 1.**
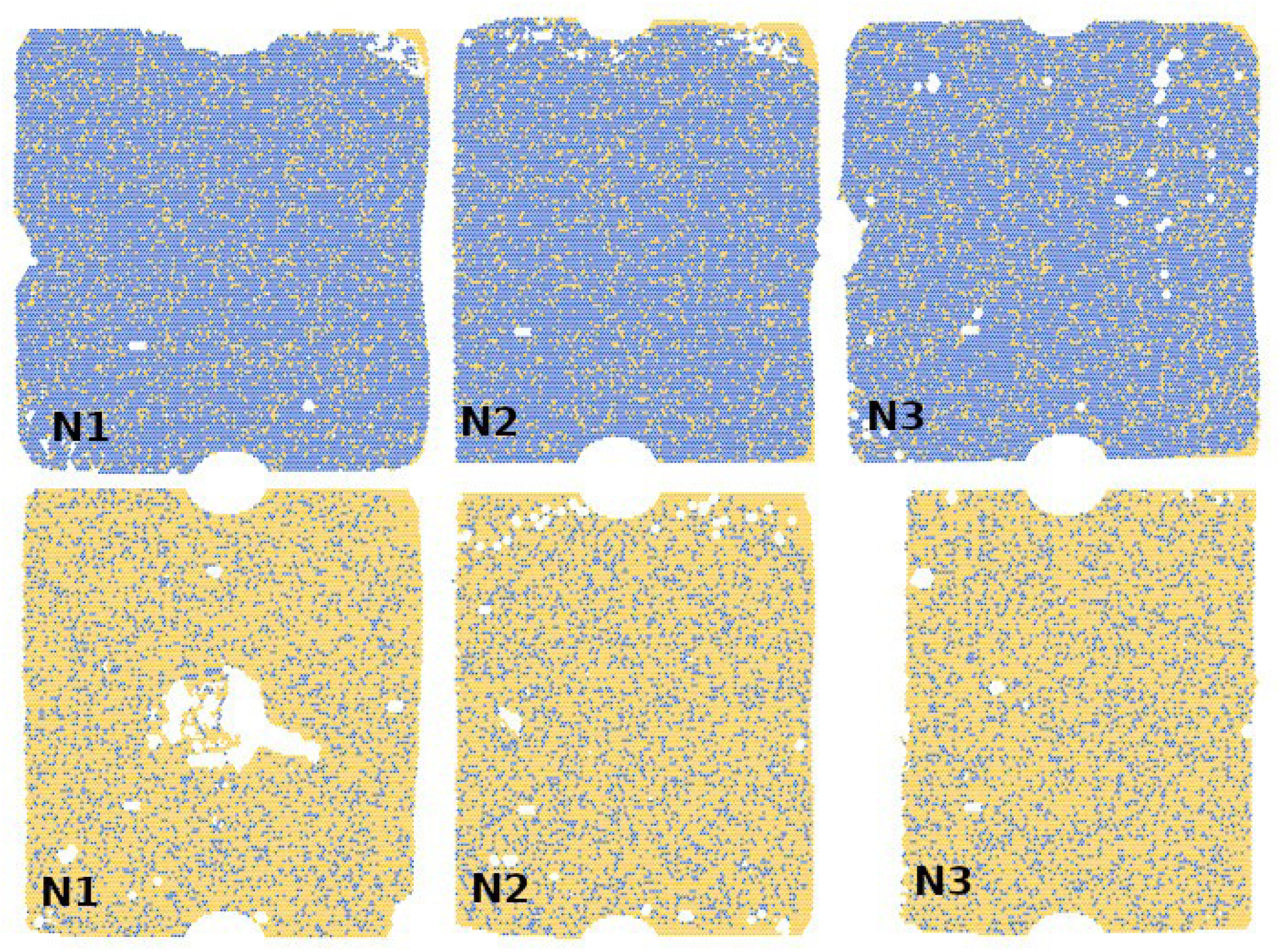
Chip images. Blue dots represent positive partitions, yellow dots represent negative partitions. Target is assigned to corresponding image, higher row: SARS-CoV2 ·10^−2^; lower row: SARS-CoV2 · 10^−3^.

Results obtained by this method can be applied in RT-qPCR reactions, using the same dilution that presented better precision values and narrower confidence interval in the quantitative curve of the RT-qPCR, and additional 10-fold dilutions above and below these values. This method is an alternative to classical method of viral titration used in RT-qPCR curves, it is faster and safer than classical methods, it can be applied in BSL2 laboratories and only cDNA is needed, therefore controls that are shared between multiple laboratories can be quantified in a single laboratory. All these situations are very common during this pandemic crisis, hence, this technique might be helpful nowadays.

